# Kaempferol suppresses the activation of mast cells by modulating the expression of FcεRI and SHIP1

**DOI:** 10.1101/2023.02.20.529186

**Authors:** Kazuki Nagata, Sanae Araumi, Daisuke Ando, Miki Ando, Naoto Ito, Yuki Ikeda, Miki Takahashi, Sakura Noguchi, Takuya Yashiro, Masakazu Hachisu, Chiharu Nishiyama

## Abstract

In the present study, we evaluated the effects of kaempferol on bone marrow-derived mast cells (BMMCs). Kaempferol treatment significantly and dose-dependently inhibited IgE-induced degranulation, and cytokine production of BMMCs under the condition that cell viability was maintained. Kaempferol downregulated the surface expression levels of FcεRI on BMMCs, but the mRNA levels of FcεRIα, β, and γ-chains were not changed by kaempferol treatment. Furthermore, the kaempferol-mediated downregulation of surface FcεRI on BMMCs was still observed in the presence of a protein-synthesis inhibitor. These results suggest that kaempferol reduced the surface FcεRI on BMMCs, independently of transcription and translation. We also found that kaempferol inhibited both LPS- and IL-33-induced IL-6 production from BMMCs, without affecting the expression levels of their receptors, TLR4 and ST2. Although kaempferol treatment increased the protein amount of NRF2, a master transcription factor of antioxidant stress, in BMMCs, inhibition of NRF2 did not alter the suppressive effect of kaempferol on degranulation. Finally, we observed that kaempferol treatment increased the levels of mRNA and protein of a phosphatase SHIP1 in BMMCs. These results indicate that kaempferol inhibited the IgE-induced activation of BMMCs by downregulating FcεRI and upregulating SHIP1, and the SHIP1 increase is involved in the suppression of various signaling-mediated stimulations of BMMCs, such as those associated with TLR4 and ST2.

## 1. Introduction

Mast cells (MCs) play important roles in allergic diseases, particularly IgE-dependent allergic responses induced by the cross-linking of the cell-type specifically expressed high affinity receptor for IgE, FcεRI, on the surface. The cross-linking of IgE-binding FcεRIs with antigen (Ag) induces the activation of MCs, resulting in the release of various mediators, such as histamine and eicosanoids, and the transactivation of cytokine genes in MCs, which are involved in rapid and late-phase allergic responses. Moreover, several polyphenols inhibit the IgE-induced activation of MCs *in vitro* or ameliorate IgE-dependent allergic responses *in vivo* [1–5]. Kaempferol is a flavonoid reported to inhibit allergic diseases in model mice [6–8]. Several *in vitro* studies using MC lines, such as rat basophilic leukemia cell line RBL-2H3 and human MC line LAD2, suggested that IgE-induced activation of MCs was suppressed by kaempferol [7,8]. However, the molecular mechanisms underlying the inhibited activation of MCs remain unclear.

In the present study, we used bone marrow-derived MC (BMMC) because we expected that BMMCs are as sensitive as natural MCs in the body to various stimulation and kaempferol, and we found that kaempferol suppressed the IgE-induced degranulation and cytokine release of BMMCs. Further analyses using flowcytometry and qPCR showed that the cell surface expression level of FcεRI was downregulated by kaempferol, which was parallel to the suppression levels of the IgE-induced activation of BMMCs and was caused in a transcription-independent manner. In addition, the activation of BMMCs by LPS or IL-33 was significantly inhibited by kaempferol, even though the expression of TLR4 and ST2 was not affected by kaempferol. We observed that kaempferol upregulated the transcription and protein expression of a phosphatase, SHIP1. Although several studies suggest that the NRF2 pathway is involved in the anti-allergic effects of natural compounds, the inhibition of NRF2 did not alter the inhibitory effects of kaempferol in the present study. Possible explanations for the downregulation of cell surface FcεRI and upregulation of SHIP1 are also discussed.

## 2. Results

### 2.1. Kaempferol suppressed IgE-mediated activation of BMMCs

To evaluate the effects of kaempferol on the activation of BMMCs, we incubated BMMCs in the presence of various concentrations of kaempferol and found that the preincubation of BMMCs with 25–50 μM of kaempferol for 24 h before the addition of IgE significantly suppressed IgE-mediated degranulation (Figure 1A), without affecting cell viability (Figure 1B). A 24 h pretreatment with 50 μM kaempferol also drastically inhibited the release of cytokines, including IL-6, TNF-α, and IL-13, from IgE-stimulated BMMCs (Figure 1C). In contrast, although the pretreatment with kaempferol tended to reduce A23187-induced degranulation of BMMCs, the effects were not significant (Figure 1D). A western blot using an anti-phospho-Tyr antibody (Ab) revealed that the IgE-induced increase of phosphorylated Tyr residues in intracellular proteins was suppressed by the kaempferol treatment (Figure 1E).

**Figure 1.**
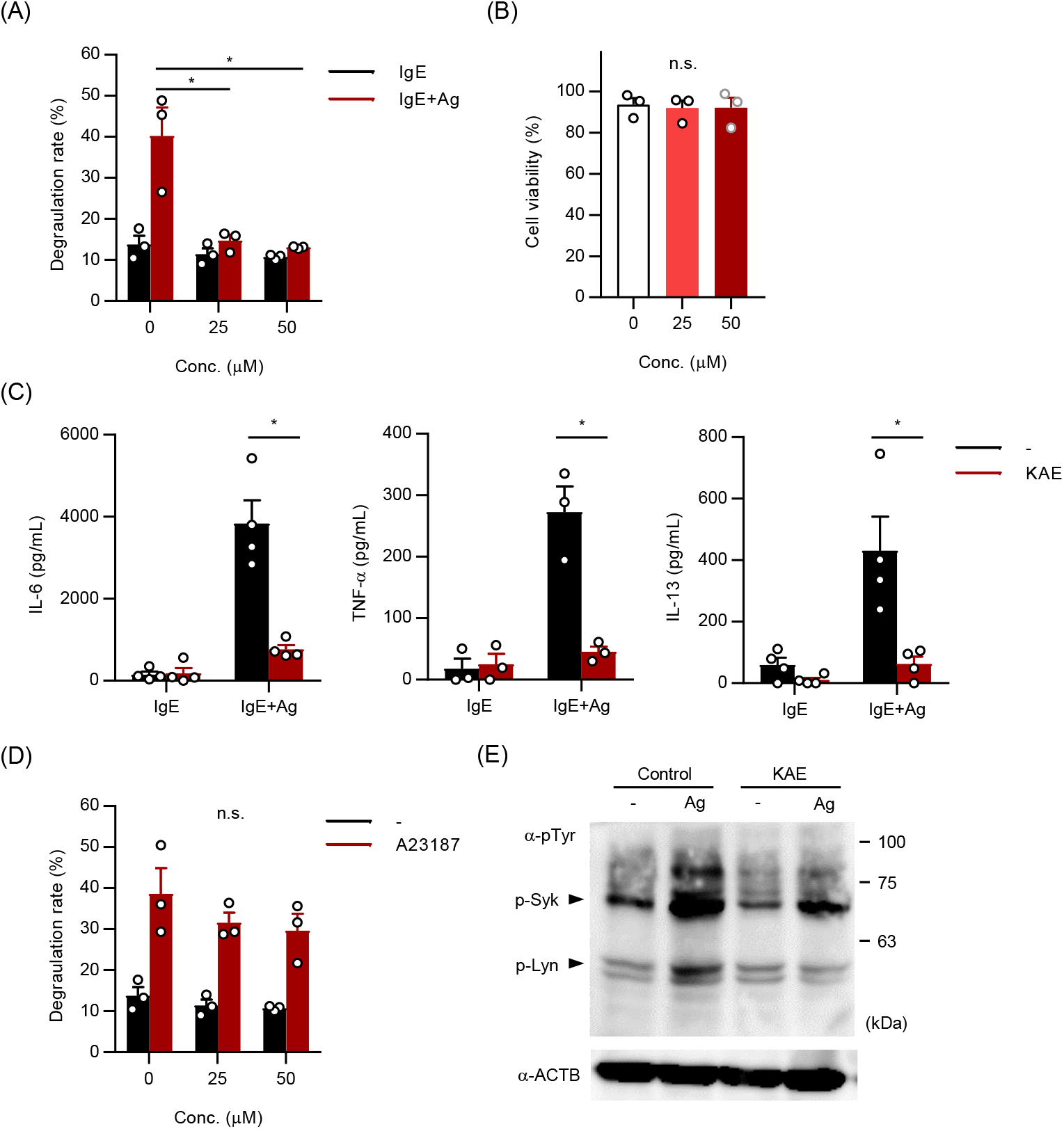
Kaempferol suppressed IgE-mediated activation of BMMCs. BMMCs were pre-incubated in the presence or absence of KAE (50 μM or indicated concentrations) for 24 h followed by each assay. (**A**) Effects of KAE on the IgE-mediated degranulation of BMMCs. After sensitization with anti-TNP-IgE, BMMCs were incubated in Tyrode’s buffer w/ or w/o TNP-BSA, and the supernatant was collected for β-hexosaminidase assay. (**B**) Effects of KAE on the viability of BMMCs. DAPI-stained cells were judged to be dead cells. (**C**) Inhibition of cytokine release from BMMCs by KAE. IgE-sensitized BMMCs were incubated in culture medium w/ or w/o TNP-BSA for 3 h and the supernatant was collected. (**D**) Effects of KAE on the degranulation of BMMCs induced by A23187. BMMCs were incubated in Tyrode’s buffer w/ or w/o A23187, and the supernatant was collected for β-hexosaminidase assay. (**E**) Western blot analysis of tyrosine-phosphorylated proteins. BMMCs were stimulated with IgE plus TNP-BSA for 10 min. The expected molecular weights of Lyn and Syk were 56 and 72 kDa respectively. The data shown in (**A-D**) represent the mean ± SEM of 3~4 independent experiments, respectively. The data shown in (**E**) represents a typical data of three other independent experiments. Dunnett’s multiple comparison test (**A**, **D**) and two tailed paired t-test (**C**) were used for statistical analyses. *, p < 0.05; n.s., not significant. Abbreviation: Conc., concentration; Ag, Antigen; KAE, Kaempferol.

These results demonstrate that kaempferol effectively suppressed IgE-mediated activation of BMMCs, mainly targeting the kinases just downstream of IgE rather than Ca^2+^-signaling.

### 2.2. Kaempferol reduced the cell surface levels of FcεRI on BMMCs

To clarify the mechanisms of the suppressive effects of kaempferol on the IgE-dependent activation of MCs, we determined the expression levels of FcεRI on kaempferol-treated BMMCs. As shown in Figure 2A, the cell surface levels of FcεRI were slightly and significantly decreased by pretreatments for 24 h with 25 and 50 μM kaempferol, respectively, under the condition that c-kit expression levels were not affected. In addition, the mRNA levels of the FcεRI subunits were not decreased, but rather increased (particularly *Fcer1a* mRNA) in the kaempferol-treated BMMCs (Figure 2B). A time course analysis showed that a significant reduction of surface FcεRI level occurred at 2 h after the addition of kaempferol in the culture medium of BMMCs, and the reduction levels were time-dependently enhanced (Figure 2C). The treatment of BMMCs with cycloheximide, an inhibitor of protein synthesis, downregulated the cell surface levels of FcεRI and the cycloheximide-induced downregulation was significantly enhanced by kaempferol (Figure 2D).

**Figure 2.**
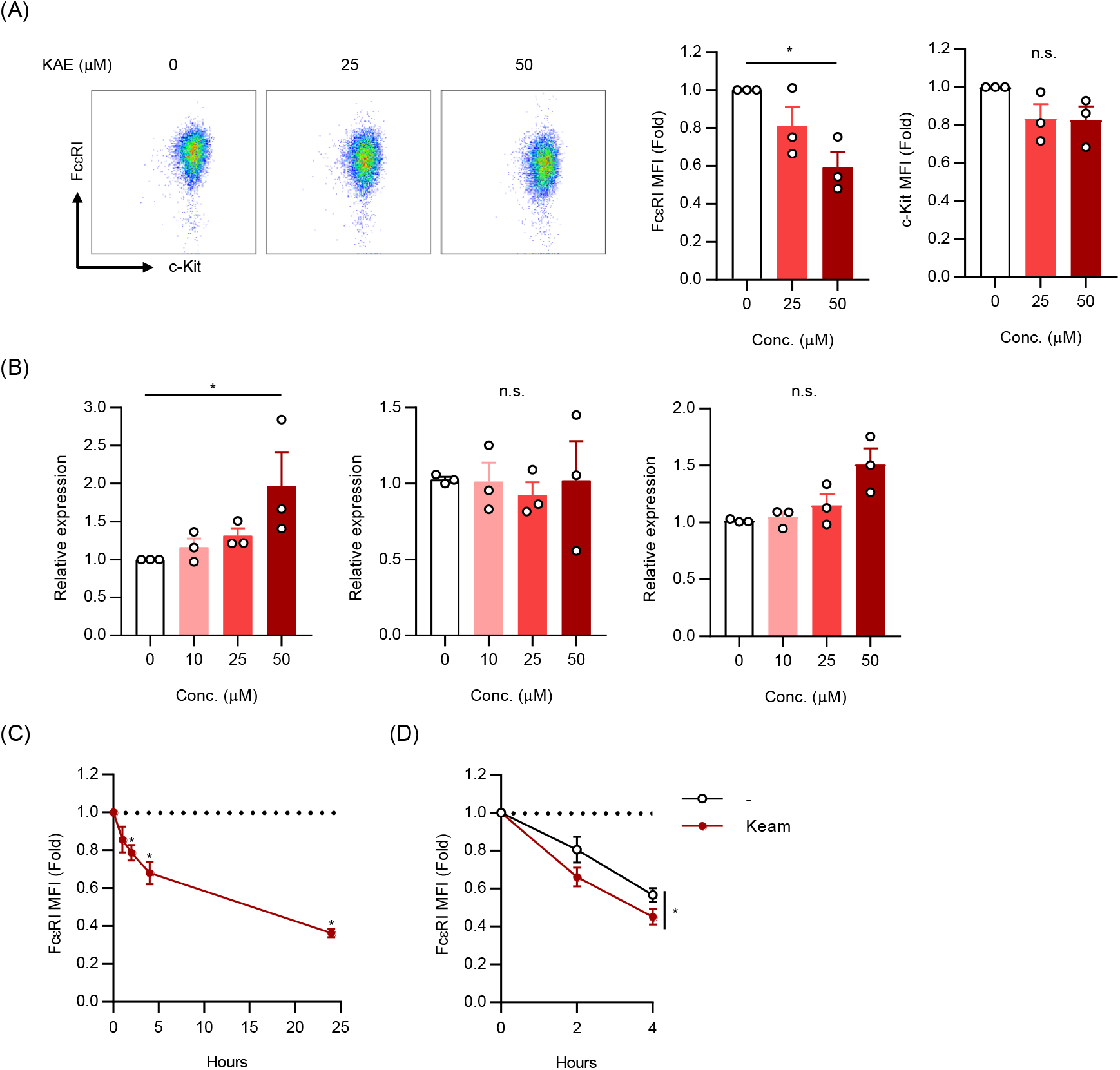
Kaempferol reduced the cell surface levels of FcεRI on BMMCs. (**A**) Cell surface expression levels of FcεRI and c-Kit. Typical dot plot profiles (left) and the mean fluorescence intensity (MFI) of FcεRI (middle) and c-Kit (right) were obtained in three independent experiments. BMMCs incubated in the presence or absence of the indicated concentrations of KAE for 24 h were stained with anti-FcεRI Ab and anti-c-Kit Ab. (**B**) mRNA expression levels of FcεRIα (*Fcer1a*), β (*Ms4a2*), and γ (*Fcer1g*) subunits. BMMCs pretreated with KAE for 24 h were harvested to assess mRNA levels by qPCR (Normalized by β-actin). (**C**) Time course of cell surface expression levels of FcεRI following KAE treatment. BMMCs were incubated in the presence of KAE (50 μM) and harvested at indicated time. (**D**) Time course of cell surface expression levels of FcεRI following KAE treatment in the presence of cycloheximide. Cycloheximide treated BMMCs were incubated in the presence or absence of KAE (50 μM) and harvested at indicated time. The data represent the mean ± SEM of 3~5 independent experiments. Dunnett’s multiple comparison test (**A**-**C**) and two tailed paired t-test (**D**) were used for statistical analyses. *, *p* < 0.05; n.s., not significant. Abbreviation: Conc., concentration; KAE, Kaempferol.

These results suggest that kaempferol suppressed cell surface expression of FcεRI in a transcription-independent manner and that the kaempferol-induced reduction of FcεRI expression was caused under the protein-synthesis inhibited condition.

### 2.3. Kaempferol suppressed LPS- and IL-33-induced IL-6 production without affecting cell surface levels of TLR4 and ST2 on BMMCs

To further clarify whether kaempferol modulates the activation of BMMCs by other stimulation, we evaluated the effects of kaempferol treatment on cytokine production from BMMCs stimulated with LPS and IL-33. The treatment with kaempferol significantly suppressed LPS-induced IL-6 release from BMMCs (Figure 3A) without exhibiting apparent effects on the cell surface expression levels of TLR4 (Figure 3B). IL-33-induced IL-6 production was also significantly reduced in kaempferol-treated BMMCs (Figure 3C), without affecting IL-33 receptor levels in BMMCs (Figure 3D).

**Figure 3.**
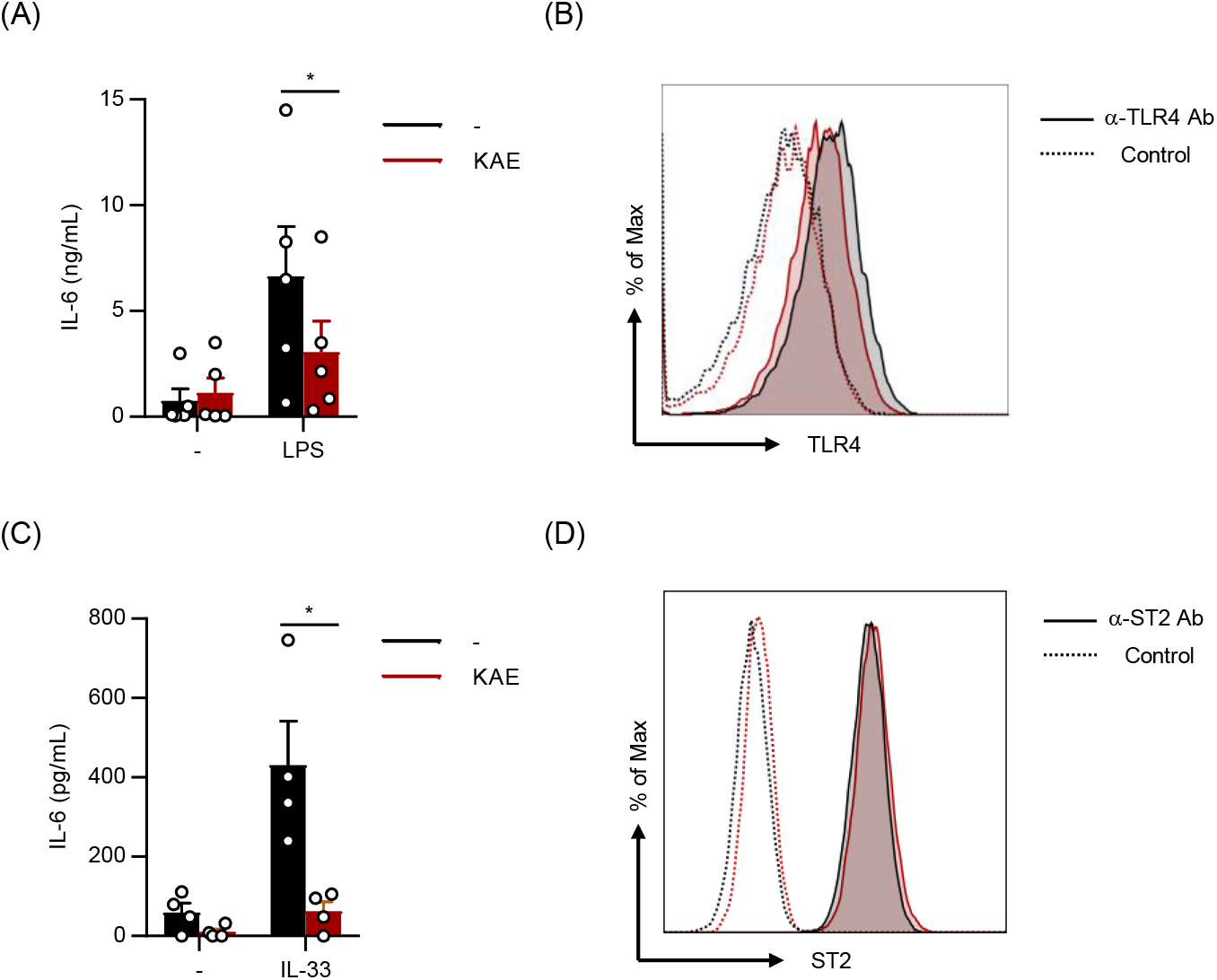
Kaempferol suppressed LPS- and IL-33-induced IL-6 production without affecting cell surface levels of TLR4 and ST2 on BMMCs. BMMCs were pre-incubated in the presence or absence of KAE (50 μM) for 24 h followed by each assay. (**A**) BMMCs were stimulated with LPS (1 μg/mL) for 3 h and the supernatant was collected to determine the concentration of IL-6. (**B**) The cell surface expression levels of TLR4 after KAE treatment. Typical histogram obtained in 2 independent experiments was shown. (**C**) BMMCs were stimulated with IL-33 (10 ng/mL) for 3 h and the supernatant was collected to determine the concentration of IL-6. (**D**) The cell surface expression levels of ST2 after KAE treatment. Typical histogram obtained in 2 independent experiments was shown. The data represent the mean ± SEM of 4 or 5 independent experiments, and two tailed paired t-test were used for statistical analyses (**A, B**). *, *p* < 0.05. Abbreviation: KAE, Kaempferol; α-TLR4 Ab, anti-TLR4 antibody; α-ST2 Ab, anti-ST2 antibody.

### 2.4. Kaempferol treatment activated NRF2 in BMMCs

The above-mentioned results showed that the activation of BMMCs by LPS and IL-33 was significantly suppressed by the kaempferol treatment under the condition that the expression of their receptors was maintained. In the case of FcεRI, cell surface expression was significantly reduced in the presence of 50 μM kaempferol, which can be involved in the suppression of IgE-induced degranulation. However, the effect of 25 μM kaempferol on FcεRI expression was not striking, but the effect on the degranulation degree was significant. Thus, we demonstrated that kaempferol modulates intercellular events induced by various stimuli.

The KEAP1-NRF2-HO-1 pathway is often activated by phytochemicals [9,10], and there are reports indicating that kaempferol activates the NRF2 pathway [11,12]. Several natural compounds, including resveratrol, exhibit anti-allergic effects by activating the NRF2 pathway in MCs [5,13–15]. Therefore, we examined the activation and role of NRF2 in kaempferol-treated BMMCs. A western blot analysis showed that the amount of NRF2 protein was increased in BMMCs 2-8 h after the addition of kaempferol (Figure 4A), suggesting that kaempferol treatment activated the NRF2 pathway in MCs. However, the IgE-mediated degranulation was reduced in kaempferol-treated BMMCs even in the presence of ML385, an inhibitor of NRF2 (Figure 4B).

**Figure 4.**
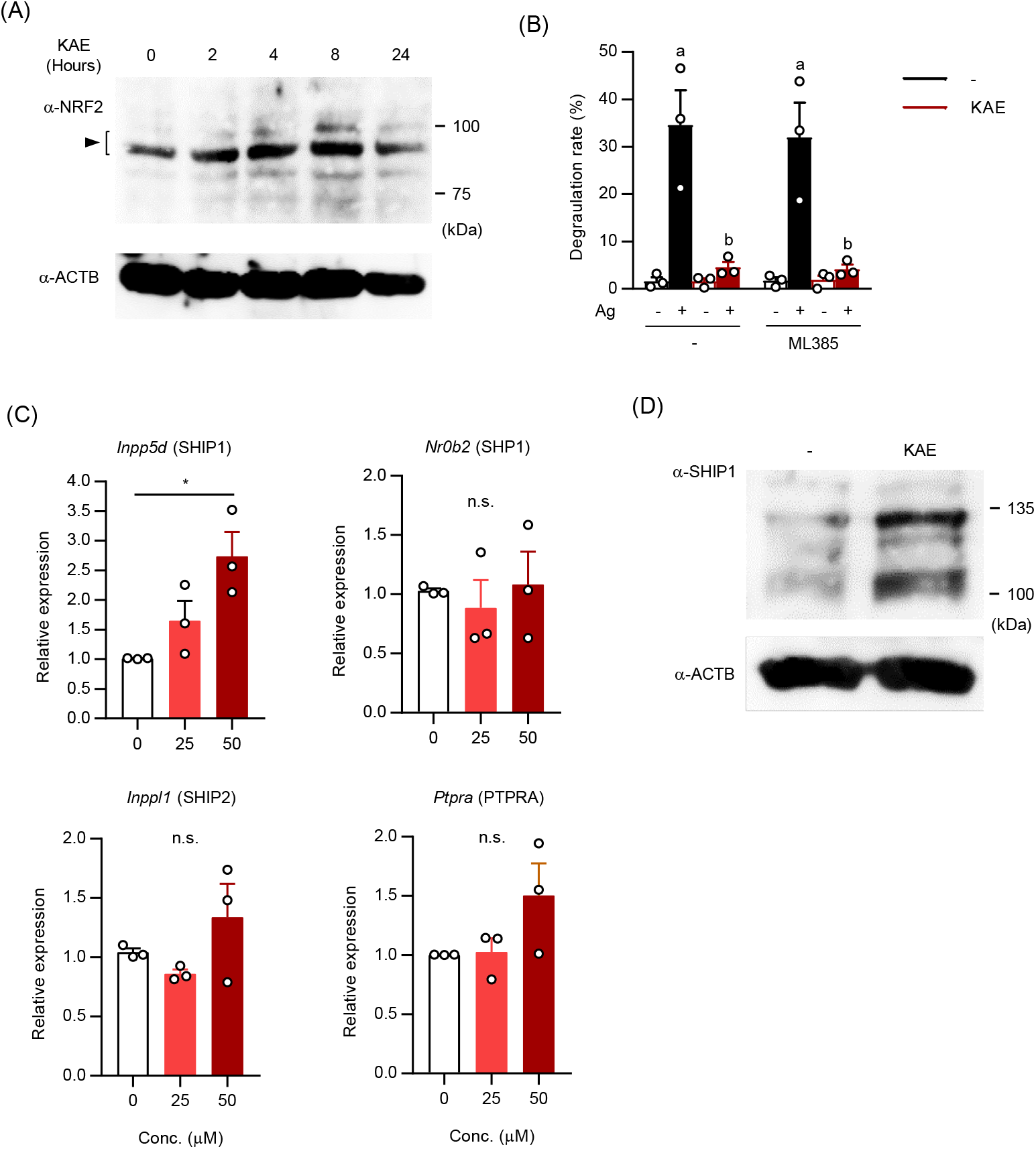
Effects of kaempferol on the NRF2 pathway and the expression of phosphatases in BMMCs. (**A**) Western blot analysis of NRF2. BMMCs were incubated in the presence of KAE (50 μM) and harvested at indicated time. The expected molecular weight of NRF2 was 97-100 kDa. (**B**) Effects of the NRF2 inhibitor on the suppression of IgE-mediated degranulation of BMMCs by KAE. BMMCs were incubated in the presence of the NRF2 inhibitor, ML385 (5 μM) and KAE (50 μM) for 24 h. Then BMMCs were sensitized with anti-TNP-IgE and incubated in Tyrode’s buffer w/ or w/o TNP-BSA. The supernatant was collected for β-hexosaminidase assay. (**C**) mRNA expression levels of SHIP1 (*Inppd5*), SHP1 (*Nr0b2*), SHIP2 (*Inppl1*), and PTPRA (*Ptpra*). BMMCs pretreated with KAE for 24 h were harvested to assess mRNA levels by qPCR (normalized by β-actin). (**D**) Western blot analysis of SHIP1 protein. BMMCs treated with 50 μM KAE for 24 h were analyzed. The data shown (**B, C**) represent the mean ± SEM of 3 independent experiments. The data shown in (**A, D**) represent a typical data of three other independent experiments. Tukey’s multiple comparison test (**B**) and Dunnett’s multiple comparison test (**C**) were used for statistical analyses. *, *p* < 0.05; n.s., not significant. Abbreviation: Conc., concentration; KAE, Kaempferol.

These results suggest that the NRF2 pathway was not involved in the suppressive effects of kaempferol on the activation of MCs, although NRF2 was activated by kaempferol treatment in MCs.

### 2.5. The levels of mRNA and protein of SHIP1 were upregulated in kaempferol-treated BMMCs

Based on the observation that the inhibitory effect of kaempferol on A23187-induced degranulation was relatively moderate compared with that on IgE-mediated degranulation, intercellular events inhibited by kaempferol were likely upstream of Ca^2+^-signaling. Considering the molecular weights of Tyr-phosphorylated proteins (Figure 1E), the kinases just downstream of FcεRI, such as Syk (72 kDa) and Lyn (56 kDa), were likely inhibited in kaempferol-treated BMMCs. Thus, we examined the effects of kaempferol on the expression of phosphatases, which inhibit the initiation of the FcεRI-mediated kinase cascade by dephosphorylating Tyr residues in ITAMs of FcεRI subunits and kinases associated with FcεRI. Among the candidate enzymes, SHIP1 was identified as a phosphatase whose mRNA level was increased in kaempferol-treated BMMCs (Figure 4C). By a western blot analysis, we confirmed that the amount of SHIP1 protein was increased by kaempferol treatment in BMMCs (Figure 4D).

From these results, we conclude that kaempferol inhibited the activation of MCs by upregulating the expression of a phosphatase SHIP1.

## 3. Discussion

Several polyphenols have been reported to possess anti-allergic activities *in vivo* and suppress the activation of MCs [1–5]. Although a flavonoid, kaempferol, was also reported to inhibit allergic responses *in vivo* in mice by modulating MC function [7,8] or other cells [6], the molecular mechanisms underlying the inhibited activation of MCs remain unclarified. In the present study, we examined the effects of kaempferol on BMMCs, which were expected to have more natural phenotypes compared with cell lines, such as RBL-2H3 and LAD2, used in the previous study [7,8]. In our study, the kaempferol treatment inhibited the activation of BMMCs upon stimulation using IgE with Ag, LPS, and IL-33. Further analyses revealed that kaempferol downregulated the cell surface expression of FcεRI and upregulated the expression of SHIP1, which may be involved in the suppressive effects of kaempferol on MC activation.

Although the cell surface levels of FcεRI were downregulated in kaempferol-treated BMMCs, the mRNA levels of all of three subunits of FcεRI were not decreased by kaempferol. Since the suppressive effects of kaempferol on FcεRI expression were still observed in the presence of cycloheximide, kaempferol appeared to downregulate the FcεRI expression in a protein synthesis-independent manner. When FcεRIα, β, and γ subunits were transiently expressed in the HEK293T cell line, surface expression of FcεRI on HEK293T cells was not reduced by the kaempferol treatment (data not shown). From these results, we suggest that kaempferol downregulated the cell surface FcεRI through a post-translational event, which is caused in MCs but not in a human kidney cell line exogenously expressing FcεRI, and endocytosis may be causing the downregulation. Although several studies have shown that IgE-binding FcεRI is internalized in the cytoplasm from the membrane and degraded, studies regarding endocytosis of empty FcεRI have not been reported. A ubiquitin ligase, CBL-B, is involved in the degradation of internalized IgE-binding FcεRI, resulting in the downregulation of cell surface FcεRI in MCs [16,17]. In our preliminary experiment, an increase in the mRNA level of CBL-B was observed in kaempferol-treated BMMCs (data not shown), although the ubiquitination of FcεRI subunits was hardly detected. To clarify this issue, further analyses are required.

SHIP1 plays inhibitory roles in IgE-induced degranulation and cytokine production of MCs because SHIP1 deficient BMMCs exhibited hyper responses in the IgE-induced degranulation [18] and IgE-induced production of proinflammatory cytokines, including IL-6 [19]. Another study revealed that SHIP1 negatively regulates TLR4-mediated LPS responses in macrophages, in which knockdown and overexpression of SHIP1 were conducted [20]. In a study regarding LPS-induced proinflammatory cytokine production from BMMCs, the amount of TNF-α released from LPS-stimulated SHIP1 knockout (KO) BMMCs was higher than that of wild-type BMMCs [21]. Considering that MC-dependent allergic inflammation was exacerbated in SHIP1 KO mice accompanied by MC hyperplasia [22], an increase of SHIP1 in MCs may exhibit anti-allergic effects *in vivo*. We also found that kaempferol effectively inhibited IL-33-induced activation of BMMCs. Although we need to clarify the detailed mechanisms of the suppressive effects of kaempferol on IL-33-mediated activation of MCs, including the involvement of SHIP1 in this effect, kaempferol may be useful to prevent or treat IL-33-mediated allergic diseases such as asthma.

In the present study, we also observed the increase of NRF2 protein levels in kaempferol-treated BMMCs. A previous study reported that resveratrol, a well-known polyphenol, suppressed C48/80-induced pseudoallergic reactions *in vivo* and MRGPRX2-mediated MC activation *in vitro* by upregulating NRF2 expression [5]. In contrast, in our experimental conditions, the inhibition of NRF2 did not affect the IgE-induced degranulation and kaempferol-mediated suppression. Although the activation of the NRF2 pathway in MCs was observed in several studies, showing the anti-allergic effects of natural compounds [5,13–15,23], the roles of NRF2 in MCs may be more complicated. In our preliminary experiments, NRF2 deficiency did not enhance IgE-induced passive anaphylaxis in mice and IgE-induced degranulation of BMMCs (data not shown). We will perform further detailed analyses of MC-dependent allergic responses of NRF2 KO mice, to reveal the roles of NRF2 in the function of MCs and the contribution of NRF2 in the anti-allergic effects of polyphenols on MCs.

Kaempferol treatment upregulated the levels of mRNA and protein of SHIP1 in MCs, suggesting the effect of kaempferol on transcription of the gene *Inpp5d*, encoding SHIP1. We will examine the roles of kaempferol on the transactivation of the *Inpp5d*. Natural compounds were often identified as ligands of nuclear receptors, and kaempferol was previously reported to function as a ligand of AhR [24]. However, in our preliminary experiments using siRNA, the involvement of AhR in the inhibitory effects of kaempferol on MCs was not observed. For other targets, we would consider miR-155 and IL-10, which are known to regulate or correlate with the expression of SHIP1 [25–28]. We should investigate whether the expressions of miR-155 and/or IL-10 are altered in kaempferol-treated BMMCs.

In the present study using BMMCs, we found that kaempferol reduced the surface expression of FcεRI and increased the expression of SHIP1, which may be involved in the inhibitory effects of kaempferol on IgE-, LPS-, and IL-33-induced activation of MCs.

## 4. Materials and Methods

### 4.1. Mice and cells

BMMCs were generated from BM cells of C57BL/6 mice (Japan SLC, Hamamatsu, Japan) by cultivation under IL-3-supplemented condition as previously described [29]. All experiments using mice were performed following the guidelines from the Institutional Review Board at Tokyo University of Science, and the present study was approved by the Animal Care and Use Committees at Tokyo University of Science: K22005, K21004, K20005, K19006.

### 4.2. Activation of MCs

A degranulation assay was performed as previously described [30,31]. Briefly, BMMCs sensitized with anti-TNP mouse IgE (clone IgE-3, BD Biosciences, San Jose, CA) were stimulated with TNP-BSA (LSL-LG1117, Cosmo Bio) in Tyrode’s buffer, and the culture supernatant was harvested to determine β-hexosaminidase activity.

To investigate the cytokine release, BMMCs were stimulated with IgE and TNP-BSA using a culture medium, instead of Tyrode’s buffer. BMMCs in culture media were also stimulated with LPS (#3024, Wako) or IL-33 (#210-33, Peprotech).

### 4.3. ELISA

The concentrations of IL-6, and TNF-α were measured using ELISA kits (ELISA MAX series, BioLegend), and of IL-13 was determined using the mouse IL-13 DuoSet ELISA kit (DY413, R&D systems).

### 4.4. Western blot analysis

Western blot analyses using anti-phospho-tyrosine (clone 4G10, Millipore), anti-NRF2 (clone D1Z9C, Cell Signaling Technology), anti-SHIP1 (clone V-19, Santa Cruz Biotechnology), and anti-β-actin (clone AC-15, Sigma-Aldrich) were performed as previously described [31].

### 4.5. Flow cytometry

Cell surface expression levels of FcεRI, c-kit, TLR4, and ST2 were determined by flow cytometric analyses using FITC-labeled anti-mouse FcεRIα (clone MAR-1, BioLegend), APC-labeled anti-mouse CD117 (clone 2B8, BioLegend), PE-labeled anti-mouse CD284 (TLR4) (clone SA15-21, BioLegend), and FITC-labeled anti-mouse T1/ST2 (clone DJ8, BioLegend), respectively. Data collected using a MACS Quant Analyzer (Miltenyi Biotech) or a FACSLyric (Beckton Dickinson) were evaluated with FlowJo software (Tomy Digital Biology, Tokyo, Japan).

### 4.6. Quantification of mRNA using real-time PCR

The total RNA extracted from BMMCs using a ReliaPrep RNA Cell Miniprep System (Promega) was reverse transcribed to cDNA with a ReverTra Ace qPCR RT Master Mix (TOYOBO, Osaka). Quantitative PCR was performed using a Step-One Real-Time PCR system (Applied Biosystems) with THUNDERBIRD SYBR qPCR Mix (TOYOBO) and primers. The primers used for measuring the mRNA levels of *Fcer1a, Ms4a2* (encoding FcεRIβ), and *Fcer1g* were described previously [31]. To measure the mRNA levels of *Inpp5d* (SHIP1), *Inppl1* (SHIP2), *Nr0b2* (SHP1), and *Ptpra* (PTPα), the following primers were used: *Inpp5d* forward; 5’-tgcagtcaatatggaacatcaag-3’, *Inpp5d* reverse; 5’-gagacgaattgagatgtgactcc-3’, *Inppl1* forward; 5’-tcagggtcactcatatcaaggtt-3’, *Inppl1* reverse; 5’-cgagtagtcttctcattccctga-3’, *Nr0b2* forward; 5’-ccacctatcgatttgaaagactg-3’, *Nr0b2* reverse; 5’-cgtttacccgagtagcgtagtaa-3’, *Ptpra* forward; 5’-cccaatactggccagaccaa-3’, *Ptpra* reverse; 5’-cctcgacagacacacggacat-3’.

### 4.7. Statistical analysis

One-way analysis of variance followed by Tukey’s multiple comparison test or Dunnett’s multiple comparison test was performed to compare three samples or more, and a two-tailed Student’s t-test was used to compare two samples.

## Abbreviations used in this paper

Ab: antibody
Ag: antigen
BMMC: bone marrow-derived mast cell
KO: knockout
MC: mast cell

